# Multiscale learning and topological analysis across complex postures enable robust nematode size quantification in pharmacological assays

**DOI:** 10.64898/2026.07.12.738008

**Authors:** Zihao (John) Li, Amanda O. Shaver, Michael E.G. Sauria, Jack Weinstein, Maya K. Mastronardo, Nikita S. Jhaveri, Kate Stone, Rachel Choo, Colin Lilley, Esha Sharma, Rohan Shrishrimal, Grayson Benson, Ariel Shi, Cecilia Soko, Erik C. Andersen

## Abstract

Body size is an important trait that reflects animal development and physiology. In nematodes, precise measurement is valuable for linking variation in body dimensions to biological questions such as developmental timing, genetic regulation, and drug responses. However, robust size measurements can be difficult to obtain because nematodes can vary in curvature, have self-intersecting postures, and overlap with neighboring animals. Current image analysis software, such as CellProfiler, can measure isolated animals in straight postures but struggles with curly or overlapping animals. Here, we present NemaSize, an artificial intelligence (AI)-aided pipeline to measure *Caenorhabditis* nematode body sizes across complex postures using multi-scale learning and topology-aware skeletonization. Using You Only Look Once (YOLO) models trained at different spatial scales, NemaSize first identifies individual animals in a large field of view (FOV; 6.75 × 6.75 mm) and then performs high-resolution body segmentation in the region of interest (ROI). Next, NemaSize converts segmented body masks into topological graph representations, allowing curly and overlapping animals to be classified and skeletonized according to the body topology. NemaSize achieved less than 4% overall error in length and width measurements across all posture classes. Compared to CellProfiler, NemaSize demonstrated higher robustness for complex postures, including a 48% error reduction in length measurements for curly or self-overlapping animals. Application of NemaSize in high-throughput imaging assays further shows that NemaSize provides accurate quantification of *Caenorhabditis briggsae* larval development in response to the anthelmintic drug ivermectin, a task that was difficult for CellProfiler because of curly animal postures. Together, NemaSize provides a robust approach for automated body size quantification for *Caenorhabditis* nematodes and will support broad applications in high-throughput pharmacological and genetic screens. Beyond nematodes, NemaSize introduces a multiscale computational framework for analyzing elongated biological objects with complex topologies.

## Introduction

Accurate body size measurement is essential for quantifying growth, development, and physiology across biological systems [1–4]. Nematodes, such as *Caenorhabditis elegans*, are powerful model organisms for these studies in part because of their simple body shape and fast life cycle [5]. In nematodes, changes in body length and width can reflect differences in growth dynamics [6,7], nutritional status [8,9], and responses to genetic or environmental perturbations [10–14]. Therefore, accurate nematode body size measurements are critical for accurate assessment of biological traits of interest and thereby improving data interpretation.

Despite the importance, body size quantification in nematodes remains technically challenging. Nematodes can adopt a wide range of complex postures, including strong curvature, self-intersection, and overlapping with surrounding animals or objects. Some methods, such as exposure to sodium azide [15–17], can be used to straighten the animals, but they can be ineffective or inapplicable in some experimental designs. These challenges limit the robustness of nematode size measurement tools, especially for those tools that are not designed to resolve complex postures [18–22]. For example, the WormToolbox of CellProfiler [19,20] can perform accurate length measurement for straight and isolated animals but struggles with curly or overlapping nematodes [23].

The challenges to resolve the complex nematode postures have been partially alleviated by the development of machine learning and computer vision, such as convolutional neural network (CNN) [24,25], Mask Regional-CNN [26], and vision transformers [27], which have enabled broad applications in behavioral tracking [28–31]. For example, a recently developed tool DeepTangleCrawl (DTC) uses a neural network trained on Tierpsy-derived [32] and hand-annotated skeletons, aided by temporal information, to maintain tracking of body pose through coils and collisions [33]. However, DTC does not extract accurate skeletons from single static images but instead requires a series of frames. The DTC-generated skeletons are not optimized for length measurement and DTC does not provide body width measurements. Similar to DTC, other AI-aided pose analysis tools are optimized for behavior tracking rather than size measurement [34–36]. Therefore, methods for body size quantification that are robust to complex postures remain underdeveloped.

To address this gap, we developed NemaSize, an AI-aided pipeline for robust nematode body size measurement across diverse postures by using multiscale learning and topology-aware skeletonization. NemaSize first uses YOLO26 models [37] trained at different spatial scales to precisely segment individual animals in microscopy images, followed by topology-based posture classification and skeletonization to measure body length and width. Validation against human ground truth showed a mean percentage error below 4% for both body length and width across posture classes. Benchmarking against the WormToolbox of CellProfiler [20] showed that NemaSize has higher robustness for complex postures (*e.g.*, 48% error reduction for curly or self-overlapping nematodes). Furthermore, we integrated NemaSize into a high-throughput larval development assay (HTLDA) [38–40] and showed that NemaSize provides accurate quantification of developmental inhibition for *C. briggsae* larvae exposed to the anthelmintic drug ivermectin. NemaSize revealed important estimators to quantify the dose-dependent drug responses, which could not be reliably obtained from CellProfiler measurements, because most animals adopted curly postures. Together, NemaSize provides a robust and scalable framework for nematode body size quantification and will support broad applications in high-throughput pharmacological and genetic screens. Beyond nematode body size measurement, NemaSize provides a multiscale strategy potentially useful for the analysis of many other elongated biological objects with complex topologies, such as neuronal processes [41–44], vascular networks [45], or plant root structures [46,47].

## Materials and methods

### Preparation of nematodes for imaging

We prepared nematodes for imaging using a HTLDA protocol [38–40]. Nematode embryos were first collected using a bleaching protocol [38], and approximately 30 embryos were then dispensed into each well of a 96-well microplate (Greiner, 655083) in 50 μL of K medium. Plates were placed in humidity chambers and incubated for 24 hours at 20°C for *C. elegans* and *C. briggsae*, and 25°C for *C. tropicalis,* with continuous shaking at 170 rpm (INFORS HT Multitron shaker). After incubation, each plate was inspected to confirm that all embryos had hatched and that animals were developmentally arrested at the first larval (L1) stage. Next, 25 μL of *E. coli* HB101 suspended in K medium was added to each well to feed the arrested L1 animals at a final concentration of OD_600_ = 10. For designated wells in the ivermectin dose response assays, the HB101 suspension was mixed with ivermectin dissolved in dimethyl sulfoxide (DMSO) to assess drug effects on larval development (see Ivermectin dose-dependent larval development assay for *C. briggsae*). Plates were incubated with continuous shaking at 180 rpm for 48 hours at 20°C for *C. elegans* and *C. briggsae*, and for 42 hours at 25°C for *C. tropicalis*. Before imaging, the animals were incubated for 10 minutes with 50 mM sodium azide in M9 solution, which immobilized the animals and allowed them to settle to the well bottom.

### Curation of annotated image dataset

Animals in 96-well plates were imaged at 2× magnification under brightfield illumination using an ImageXpress Nano system (Molecular Devices, San Jose, CA) (Fig. 1A). Images were captured using an objective lens positioned underneath the plate and recorded with a 4.7-megapixel CMOS camera. Data were saved as 2,048 × 2,048-pixel 16-bit TIFF files. Each image had a 6.75 × 6.75 mm FOV, which covered the entire well (Fig. 1A, middle). In this study, we refer to the raw image capturing the entire well as a full-well image. After data acquisition, we manually selected 227 representative full-well images containing animals with diverse postures, such as straight, curly, or overlapping animals (Fig. 1B and Table S1). The selected images also captured variation in image quality, such as the presence of background debris, poor contrast, or close proximity to the well edge (Fig. 1B). Approximately 80% of the selected images were from *C. elegans*, 10% from *C. briggsae*, and 10% from *C. tropicalis.* The selected images were uploaded to Roboflow [48], an online web-based tool for image annotation. In Roboflow, individual animals were manually annotated by outlining their body contours to generate segmentation masks for each nematode (Fig. 1A, right). For animals partially occluded by surrounding worms or other objects, such as the mutually overlapping worms (Fig. 1B), human annotators interpolated the occluded regions and drew a continuous contour representing the entire animal. The annotations were reviewed to ensure accurate delineation of animal boundaries. To standardized manual posture annotation, we defined a set of descriptors that captured common variations in nematode postures (Table S1). For each annotated animal, a human assigned a posture class using the predefined posture descriptors (Tables S1 and S2). The final dataset contained 7,960 annotated animals spanning 16 posture classes (Table S2). After annotation, the dataset was exported in a YOLO-compatible format containing the original images and corresponding mask-based labels, which defined the posture class and pixel-level location of each animal within the image (Fig. 1A, right). This annotated dataset was subsequently used to train and evaluate the YOLO models for animal identification and segmentation.

**Figure 1.**
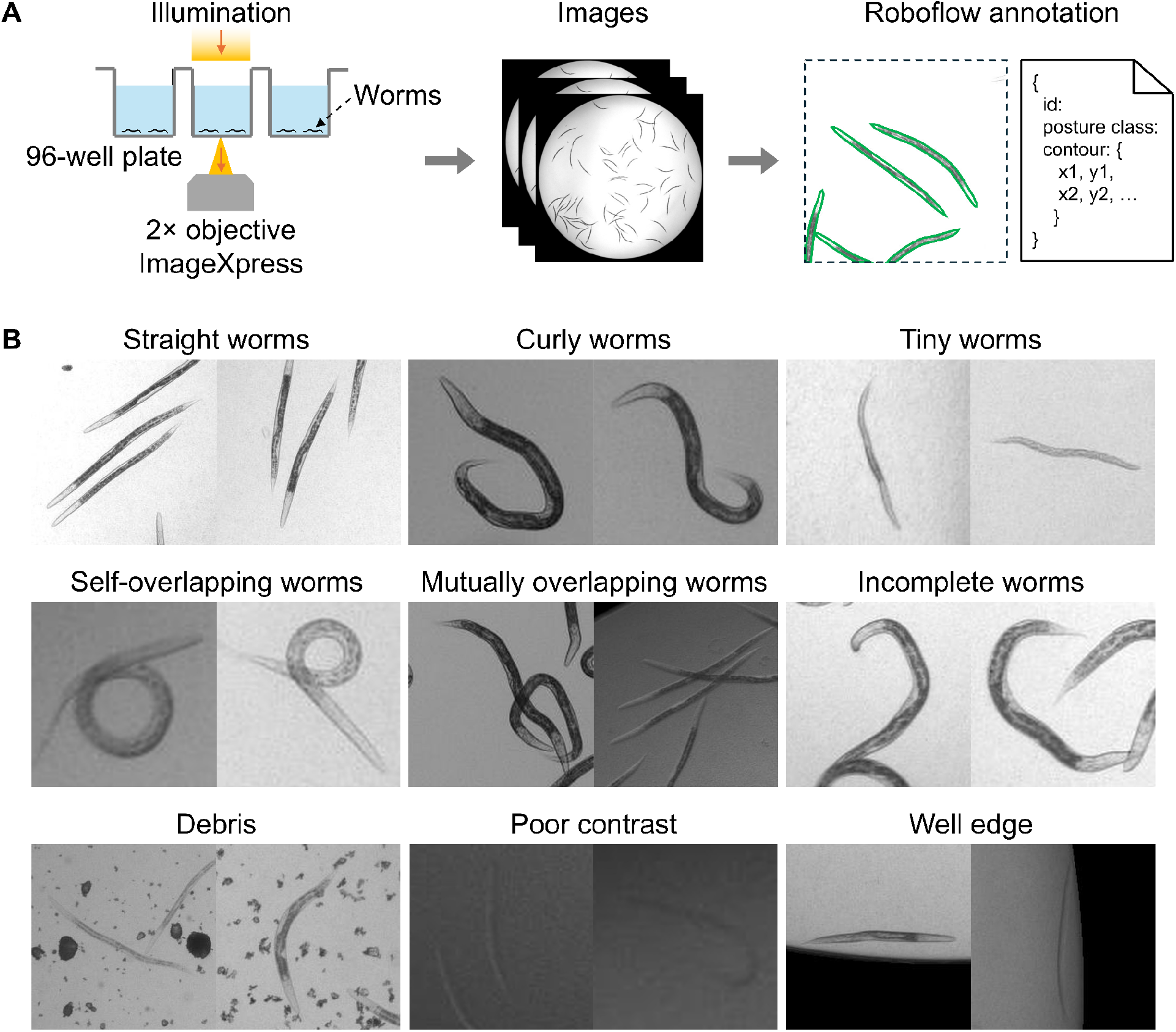
Overview of image dataset for training of YOLO26-WF and WS. (A) Workflow of image acquisition and data annotation. Images of nematodes on 96-well plates were acquired using a motorized 2× objective lens in the Molecular Devices ImageXpress system. Contours for individual animals (green solid lines) were manually annotated using the Roboflow software (see Curation of annotated image dataset). FOV = 6.75 × 6.75 mm (middle) and 1.26 × 1.26 mm (right). (B) Representative ROI images in the dataset showing diverse body postures and image quality used for training YOLO26-WF and WS. Square FOV side lengths are 790 µm (left) and 748 µm (right) for Straight worms; 389 µm (left) and 405 µm (right) for Curly worms; 405 µm (left) and 478 µm (right) for Tiny worms; 270 µm (left) and 306 µm (right) for Self-overlapping worms; 550 µm (left) and 820 µm (right) for Mutually overlapping worms; 461 µm (left) and 389 µm (right) for Incomplete worms; 810 µm (left) and 662 µm (right) for Debris; 329 µm (left) and 227 µm (right) for Poor contrast; and 632 µm (left) and 655 µm (right) for Well edge.

### Training of YOLO26 models

We trained two YOLO26x-seg models [37,49] using the Ultralytics library in Python 3.12.12 (Fig. 2). YOLO, short for “You Only Look Once”, is a family of CNN-based deep learning models that predict object locations, segmentation masks, and confidence scores from each input image through a single-stage forward inference [24,50].

**Figure 2.**
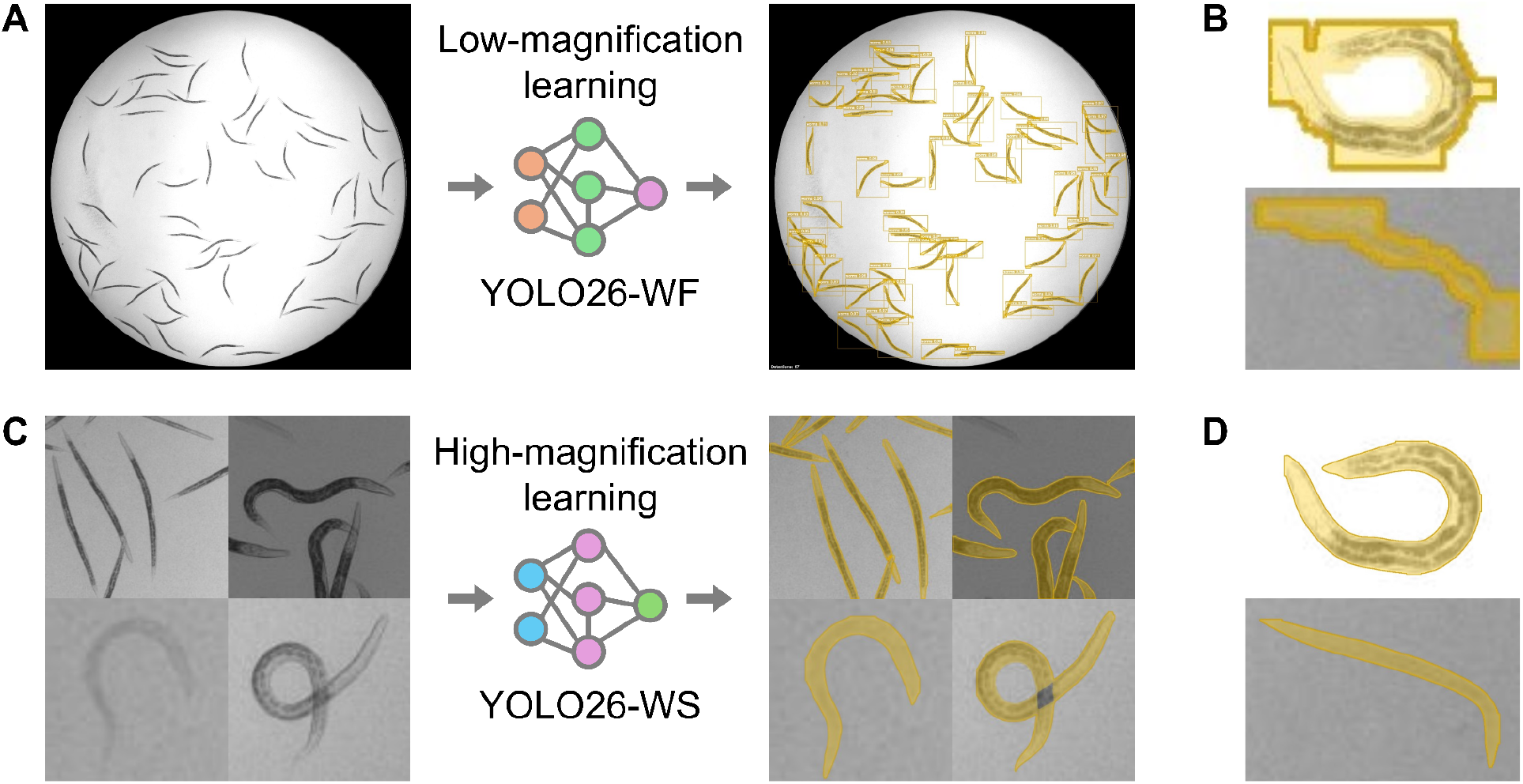
Multiscale learning for robust identification and segmentation of individual *Caenorhabditis* animals across different body postures. (A) A YOLO26-WF model was trained on full-well images for animal identification. FOV = 6.75 × 6.75 mm. (B) Representative animals showing suboptimal segmentation by YOLO26-WF, where predicted masks (yellow masks) inaccurately followed the body boundaries. FOV = 319 × 255 µm (upper) and 306 × 204 µm (lower). (C) A YOLO26-WS model was trained on ROI images showing individual animals with different postures. FOV = 965 × 965 µm (upper left), 524 × 524 µm (upper right), 191 × 191 µm (lower left), and 296 × 296 µm (lower right). (D) Improved segmentation by YOLO26-WS for the same animals shown in (B).

The first model, YOLO26-Worm Finding (YOLO26-WF), was trained for animal identification in full-well images (Fig. 2A and B). YOLO26-WF was initialized using the official YOLO26x-seg weights pretrained on the Common Objects in Context (COCO) dataset [51]. The dataset contained 227 full-well images with 182 (80%) used for training and 45 (20%) for validation. Although the animals were annotated with different posture classes (see Curation of annotated image dataset), all annotations were merged into a single “worm” class before training. The model was trained for 100 epochs using full-well images resized from 2,048 × 2,048 to 928 × 928 pixels, with a batch size of eight.

The second model, YOLO26-Worm Segmentation (YOLO26-WS), was trained for body segmentation (Fig. 2C and D). Different from YOLO26-WF, which was trained on full-well images, YOLO26-WS was trained on ROI images of nematodes. For each annotated worm, a square ROI was created centered at the animal annotation. The side length of the ROI, *S*, was set to 1.2 times the larger dimension of the annotation bounding box, *i.e*., *S* = 1.2 × max(width, height), which provided a margin around the animal to ensure that the entire body was captured (Fig. 2C). The corresponding region was then cropped from the original full-well image, and the cropped image was resized to 640 × 640 pixels. In this study, we refer to such a cropped full-well image centered on an animal of interest as an ROI image.

The training and validation ROI image datasets for YOLO26-WS were generated from the corresponding full-well training and validation images used for YOLO26-WF, thereby preserving the same data split. The ROI cropping generated 6,428 training images and 1,532 validation images for YOLO26-WS. Transfer learning was then performed by initializing YOLO26-WS with the best checkpoint from the YOLO26-WF training. YOLO26-WS was trained on the ROI images for 200 epochs with a batch size of 16. Both YOLO26-WF and YOLO26-WS were trained on an NVIDIA GeForce RTX 3090 graphical processing unit (GPU) with 24 GB of video random access memory (VRAM). Stochastic gradient descent was used as the optimizer, with an initial learning rate of 0.01, momentum of 0.937, and weight decay of 0.0005. Data augmentations were applied during training, including random brightness variation, translation, scaling, horizontal flipping, and mosaic augmentation. Model checkpoints were saved every 10 epochs. For each model, the best checkpoint was selected based on the validation performance. The best checkpoint was used for subsequent animal identification or segmentation tasks.

### Performance metrics for nematode identification and segmentation

Model performance was evaluated on the validation set using standard object detection and instance segmentation metrics including precision, recall, mAP50, and mAP50-95 (Table 1) [52]. For animal identification, predicted bounding boxes were compared with manually annotated bounding boxes. For segmentation, predicted masks were compared with manually annotated worm masks. A prediction was considered a true positive when its intersection over union (IoU) with the corresponding ground-truth annotation was equal to or greater than 0.50. Precision was calculated as TP/(TP + FP), where TP is the number of true positive detections, and FP is the number of false positive detections. Recall was calculated as TP/(TP + FN), where FN is the number of missed ground-truth annotations. Mean average precision at IoU 0.50 (mAP50) was calculated as the area under the precision-recall curve using an IoU threshold of 0.50. mAP50-95 was calculated by averaging AP values across IoU thresholds from 0.50 to 0.95 in an increment of 0.05. Thus, mAP50-95 provides a stricter evaluation of identification or segmentation performance compared to mAP50.

**Table 1.**
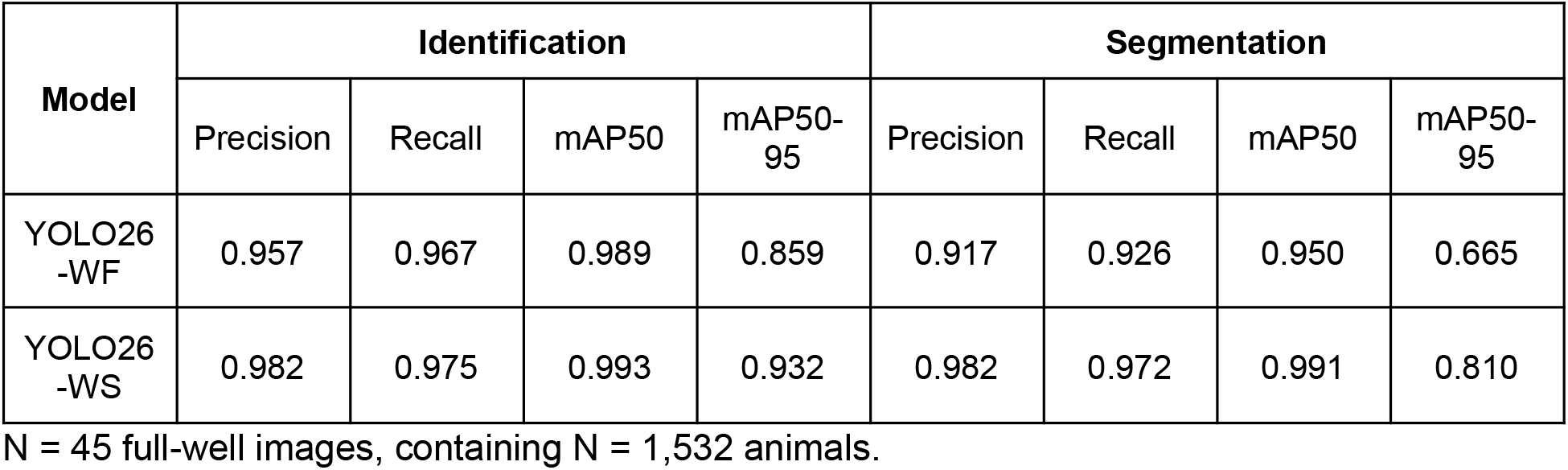
Performance metrics of nematode identification and segmentation.

| **Model** | **Identification** |  |  |  | **Segmentation** |  |  |  |
| --- | --- | --- | --- | --- | --- | --- | --- | --- |
|  | Precision | Recall | mAP50 | mAP50-95 | Precision | Recall | mAP50 | mAP50-95 |
| YOLO26-WF | 0.957 | 0.967 | 0.989 | 0.859 | 0.917 | 0.926 | 0.950 | 0.665 |
| YOLO26-WS | 0.982 | 0.975 | 0.993 | 0.932 | 0.982 | 0.972 | 0.991 | 0.810 |

N = 45 full-well images, containing N = 1,532 animals.

### Dual-network inference for nematode segmentation

Animal segmentation was performed using a dual-network inference workflow. First, the YOLO26-WF model was applied to full-well images to identify individual animals and generate bounding boxes around each detected animal. For each detection, a square ROI was cropped from the original full-well image, with the side length, *S*, set to 1.3 times the larger dimension of the bounding box, i.e. *S* = 1.3 × max(width, height), to ensure that the ROI captured the entire animal body. This ROI cropping step matched the input format used to train YOLO26-WS (see Training of YOLO26 models). Next, each ROI image was analyzed by the YOLO26-WS model to generate pixel-wise segmentation masks for the detected animals. Metadata for each ROI, including its size and coordinates relative to the original full-well image, was stored in JSON format. These metadata were later used to map the resulting segmentation masks back to the coordinate system of the original full-well image in the downstream analyses.

### Graphical analysis and topological classification

After pixel-wise animal segmentation by YOLO26-WS, each segmentation mask was converted into a binary mask for topology classification (Fig. 3). Mask preprocessing was performed to reduce segmentation artifacts. For example, small internal holes and local discontinuities caused by segmentation artifacts were filled. The mask contour was also smoothed to reduce sharp corner artifacts introduced by the polygon-based YOLO segmentation. Next, a medial axis was computed from the preprocessed binary mask (Fig. 3A) [53,54]. The medial axis represents the set of points within the animal body that are locally equidistant from the mask boundary, thereby, providing a representation of the body centerline. The medial axis was then converted into a graph representation, in which skeleton branches were represented as edges and branch endpoints or junctions were represented as nodes (Fig. 3A). Animal topology was then classified based on the graph structure (Fig. 3A).

**Figure 3.**
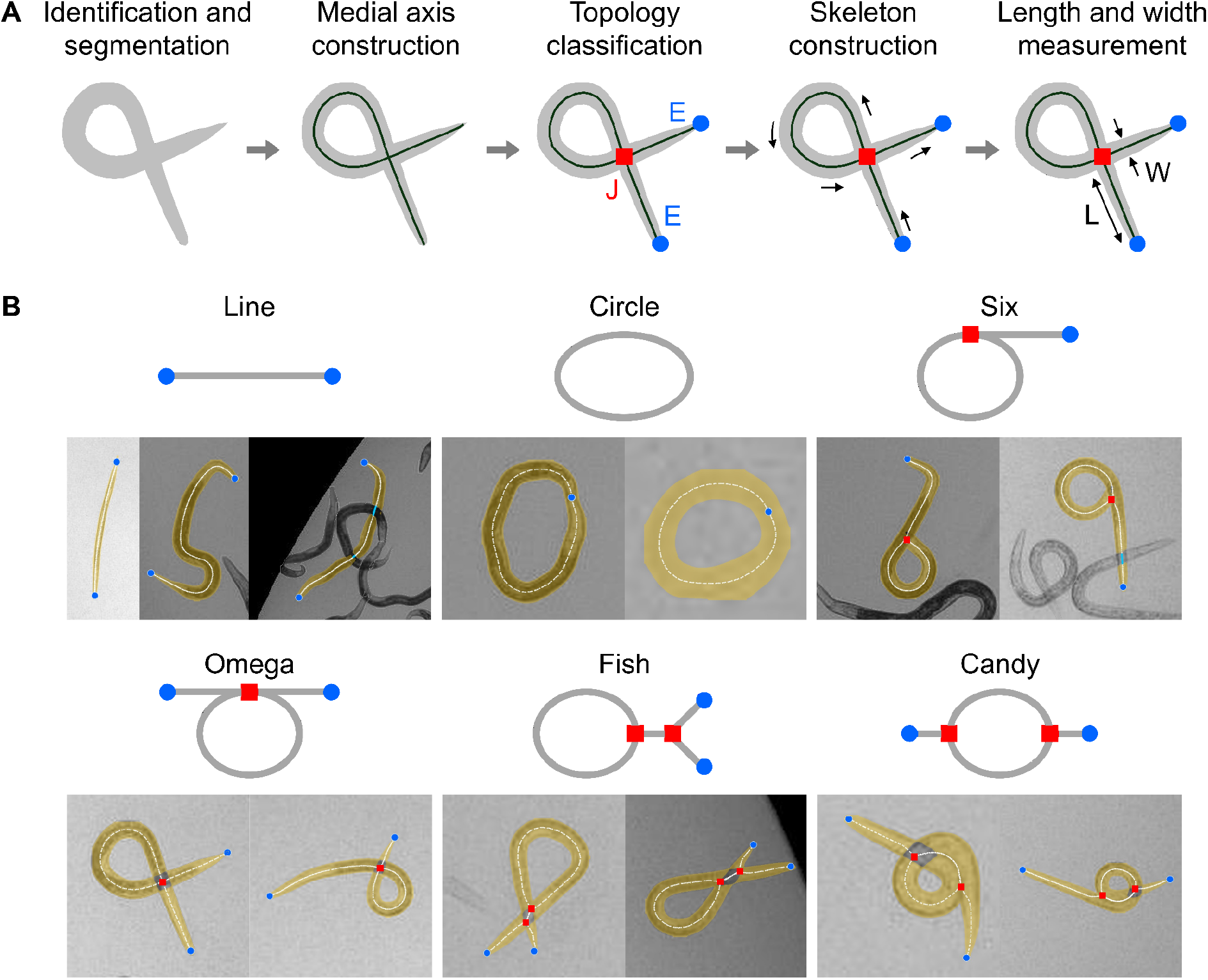
Robust skeletonization and body size measurements across different postures are enabled by topological analyses. (A) Schematics of the topology-aware skeletonization and body size measurement. J: junction; E: end point; L: length; and W: width. (B) Representative results of skeletonization across different posture topologies (topology definition see Graphical analysis and topological classification). Yellow masks denote segmented body masks, red squares denote junctions, blue circles denote object end points, and white lines denote constructed skeletons. FOV = 380 × 949 µm (left), 318 × 530 µm (middle), and 652 × 652 µm (right) for Line; 343 × 343 µm (left) and 105 × 105 µm (right) for Circle; 458 × 458 µm (left) and 418 × 418 µm (right) for Six; 372 × 372 µm (left) and 352 × 352 µm (right) for Omega; 306 × 306 µm (left) and 356 × 356 µm (right) for Fish; and 231 × 231 µm (left) and 422 × 422 µm (right) for Candy.

NemaSize classifies the graph structure of the medial axis into six topology classes according to the following definitions (Fig. 3B). The Line class is defined as an acyclic graph with two free endpoints. The Circle class is defined as a closed loop with no free endpoints. The Six class is defined as a graph containing one self loop and one free endpoint. The Omega and Fish classes both contain two free endpoints and one self loop, and are distinguished by the number of junction nodes connecting the self loop to the endpoints: Omega contains one junction node, whereas Fish contains two junction nodes. Finally, the Candy class is defined as a cyclic graph in which two distinct branches connect the same pair of junction nodes, forming a non-self-loop cycle. Each junction node is also connected to one free endpoint through a terminal branch. These topology classifications were used to guide subsequent skeleton construction.

### Topology-aware skeleton construction

After topology classification, NemaSize constructed a skeleton for each animal based on its medial-axis graph structure (Fig. 3B). The goal of skeleton construction was to generate a single ordered path from one body end to the other. Therefore, any graphs containing a loop were converted into an ordered skeleton sequence by assigning start and end points along the loop according to the corresponding topology.

For Line animals, the topological graph was acyclic, and the skeleton was generated by tracing the medial axis between the two endpoints.

For Circle animals, which contained a self loop without free endpoints, a start/end point was assigned at the narrowest point of the body mask, thereby converting the closed loop into an ordered skeleton sequence.

For Six animals, the skeleton sequence started at the free endpoint of the terminal branch, extended through the branch-loop junction, and then traversed the self loop back to the same junction, which defined the end of the ordered skeleton sequence.

For Omega and Fish animals, the medial-axis graph contained two free endpoints and one self loop. For both classes, the skeleton sequence was constructed by traversing from one endpoint branch into the loop and exiting the loop through the other endpoint branch. Because the self loop could be traversed in two possible directions, NemaSize evaluated both directions and selected the one that produced the smoothest transition at the branch-loop junctions, based on the local tangent direction of the skeleton.

For Candy animals, the graphs contained a non-self-loop cycle formed by two distinct branches connecting the same pair of junction nodes. NemaSize evaluated candidate paths in which one branch was traversed twice to represent the self-overlapping body segment, whereas the other branch was traversed once. The candidate skeleton with the smoothest transitions at the junction nodes was selected.

After the topology-specific skeleton was assembled, the ordered skeleton coordinates were smoothed using a B-spline [55,56] and resampled to 200 points to generate the final skeleton representation for downstream body size measurements.

### Occlusion handling

Occlusion by neighboring animals or other objects can cause a single animal mask inferred by YOLO26-WS to appear as multiple disconnected fragments in the ROI image (Fig. 3B). To reconstruct a skeleton for the complete animal, NemaSize bridges gaps between fragments when the connection is geometrically plausible. NemaSize first discarded components with areas smaller than 200 pixels² in the ROI image, because they were likely caused by segmentation artifacts. Next, individual fragments were analyzed independently to obtain a skeleton line for each fragment. For each gap, the algorithm calculated the length of the bridge *D*, and the angles *θ* between the bridge and the fragment skeletons at the connection points. A bridge was accepted only when *D* was less than 110 pixels and each angle *A* was less than 110°. If no candidate bridge was accepted, the skeleton of the mask fragment closest to the ROI image center was retained as the fallback skeleton.

### Measurement of body length and width

After the skeleton was generated in the ROI image, body length *L* was measured as the total arc length of the skeleton line, calculated as the sum of the Euclidean distances between each pair of adjacent skeleton points. The average body width *W* was calculated as the segmented body area *A* divided by *L*. For animals with fragmented body masks because of partial occlusion, the interpolated skeleton segments spanning the gaps between fragments (see Occlusion handling) were excluded from the calculation of *L*, to prevent underestimation of *W*. Next, the length and width measured in the ROI image were scaled back to the coordinate system of the original full-well image using the metadata generated previously by the dual-network segmentation workflow (see Dual-network inference for nematode segmentation). Pixel-based measurements were also converted to micrometers using the calibrated magnification factor of 3.29 µm per pixel in the full-well image.

### Performance comparison to CellProfiler

To evaluate NemaSize performance, we compared NemaSize measurements with measurements from the WormToolbox of CellProfiler [19,20] and human annotations using the validation dataset (N = 45 full-well images). The human-annotated masks and their corresponding body size measurements were used as the ground-truth data. We first used the CellProfiler software (Version 4.0.3) [19,20] to measure nematode body length in the validation image set. A Nextflow pipeline (version 24.10.1) (https://github.com/AndersenLab/cellprofiler-nf) was written to run command-line instances of CellProfiler in parallel on the High-Performance Computing (HPC) Cluster. The custom CellProfiler pipeline generated animal length measurements using four worm models: three worm models tailored to capture animals at the L4 larval stage, in the L2 and L3 larval stages, and the L1 larval stage, respectively, as well as a “multi-drug high dose” (MDHD) model, to capture animals with more abnormal body morphologies caused by extreme drug responses [40,57]. The worm model estimates and custom CellProfiler pipelines were written using the WormToolbox in the graphical user interface (GUI)-based instance of CellProfiler [58]. Next, a custom R package, easyXpress (Version 2.0), was used to process animal measurements output from CellProfiler [23,59]. We used easyXpress to filter the measurements from worm objects with problematic model fitting, close proximity to the well edge, or statistical outliers within individual wells.

Animals identified by NemaSize and CellProfiler were independently matched to manually labeled animals using IoU. NemaSize matches were based on mask IoU with a threshold of 0.70, whereas CellProfiler matches were based on bounding-box IoU with a threshold of 0.30. For each manually labeled animal, the computer-detected object with the highest IoU was selected as the match if its IoU exceeded the corresponding threshold. Otherwise, the animal was considered unmatched. The matching workflow ensured that the same animals were analyzed by both tools and correctly compared with the corresponding human annotations. For each matched animal, body length values from NemaSize and CellProfiler were compared with the corresponding human measurements (Fig. 4A). Percentage error was calculated as the difference between the computer and human measurements, divided by the human measurement. Percentage errors were calculated across different posture categories (Fig. 4B).

**Figure 4.**
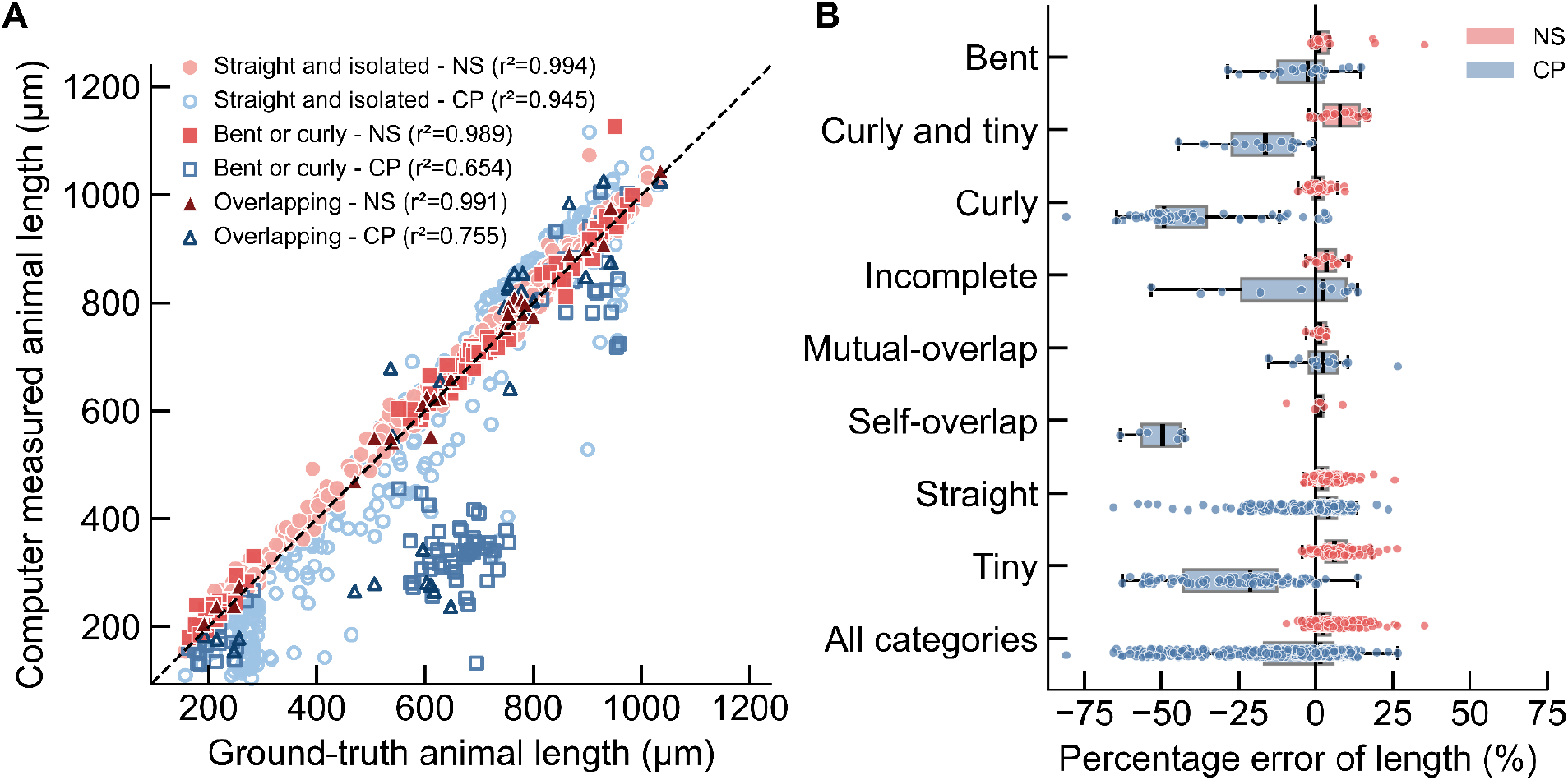
Measurements of body length from NemaSize (NS) and CellProfiler (CP) compared to manually measured truth sets. (A) A scatter plot showing body length measured by NemaSize and CellProfiler versus ground-truth manual measurements. Each point represents a single animal. N = 518 animals for straight and isolated worms, N = 92 for bent or curly worms, and N = 29 for overlapping worms. (B) A box plot showing the percentage error of body length measurements for NemaSize and CellProfiler as compared to manual measurements across different posture classes (Tables S1 and S2). Each point represents a single animal. N = 16 animals for bent worms, N = 16 for curly and tiny worms, N = 60 for curly worms, N = 11 for incomplete worms, N = 12 for mutually overlapping worms, N = 6 for self-overlapping worms, N = 423 for straight worms, N = 95 for tiny worms, and total N = 639 for all categories. Data are shown as Tukey box plots with the median as a solid black vertical line, the left and right edges of the box representing the 25^th^ and 75^th^ percentiles, respectively. Whiskers extend to the most extreme data point within 1.5 times the interquartile range from the box edges.

### Grouping of posture classes for data visualization and analysis

In the original annotated dataset, each animal was assigned to one of the 16 predefined posture classes (see Curation of annotated image dataset; Tables S1 and S2). To facilitate data visualization and analysis, some posture classes were grouped according to the rationale described below. For the scatter plot showing the consistency between NemaSize and ground-truth measurements, we merged the original posture classes to reduce the number of data groups and improve the readability of the makers (Fig. 4A). The “Straight” and “Tiny” classes were merged into the “Straight and isolated” class (Tables S1 and S2; Fig. 4A). Any original posture class containing the descriptor “Bent” or “Curly” was grouped into the “Bent or curly” class (Tables S1 and S2; Fig. 4A). Finally, any original posture class containing the descriptor “Mutually overlapping”, “Self-overlapping”, or “Incomplete” was merged into the “Overlapping” class (Tables S1 and S2; Fig. 4A). For quantification of NemaSize measurement error, original posture classes with fewer than five animals were merged into the most similar posture class to maintain sufficient sample sizes for statistical analysis (Figs. 4B and S1). The “Incomplete and curly” class (N = 2 animals) and “Incomplete and tiny” class (N = 4 animals) were both merged to the “Incomplete” class (Table S2; Figs. 4B and S1). The “Tiny and bent” class (N = 4 animals) was merged into the “Bent” class (Table S2; Figs. 4B and S1).

### Ivermectin dose-dependent larval development assay for *C. briggsae*

We followed the HTLDA protocol [38–40] to assess *C. briggsae* larval development in response to different doses of ivermectin. A 100 µM ivermectin stock solution (Sigma-Aldrich, Catalog # I8898-1G) was prepared in DMSO (Fisher Scientific, Catalog # D128-1), aliquoted, and stored at -20°C until used. The ivermectin stock solution was freshly thawed at room temperature and diluted to working concentrations right before the experiments. After bleach synchronization (see Preparation of nematodes for imaging), the arrested L1 animals were fed an *E. coli* HB101 suspension mixed with ivermectin working solution. The animals were exposed to HB101 at a final concentration of OD_600_ = 10 and ivermectin at one of the 12 concentrations, including 0, 0.029, 0.058, 0.11, 0.23, 0.46, 0.93, 1.8, 3.7, 7.5, 15, and 30 nM, in a final volume of 75 μL per well. Next, the animals were grown for 48 hours at 20°C with continuous shaking at 180 rpm before imaging on the ImageXpress platform. Before image acquisition, the animals were immobilized using sodium azide in M9 solution (see Preparation of nematodes for imaging).

The effects of ivermectin on *C. briggsae* larval development were assessed by measuring animal body length in images acquired on the ImageXpress platform (Fig. 5). To quantify the extent to which ivermectin inhibited larval development, we calculated normalized animal length for each *C. briggsae* strain at each dose. The normalized animal length was calculated as the difference between the median body length measured at a given ivermectin dose and the median body length measured in the zero-dose control condition for the same strain (Fig. 5D and E). A more negative normalized animal length indicated stronger inhibition of larval development.

**Figure 5.**
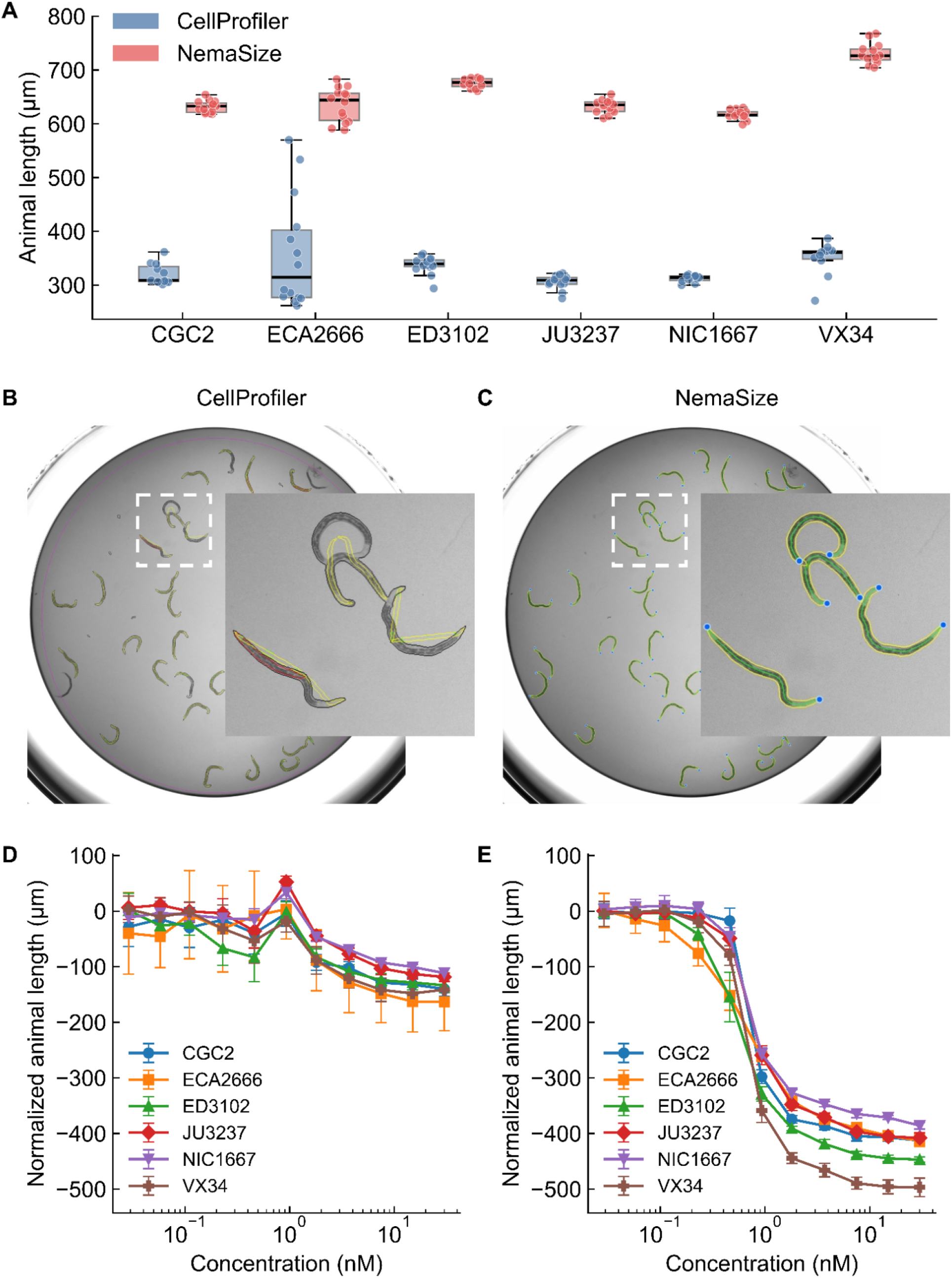
NemaSize captures accurate animal lengths and improves the interpretation of HTLDA. (A) Median animal length values from populations of six *C. briggsae* wild strains grown in DMSO are shown on the y-axis. Blue: CellProfiler; Red: NemaSize. Data are shown as Tukey box plots with the median as a solid horizontal line, the top and bottom of the box representing the 75^th^ and 25^th^ quartiles, respectively. The top whisker is extended to the maximum point that is within a 1.5 interquartile range from the 25^th^ quartile. Each point represents a single well. (B and C) A representative full-well image (FOV = 6.75 × 6.75 mm) of VX34 analyzed by (B) CellProfiler and (C) NemaSize. The region enclosed by the white dashed line (FOV = 1.29 × 1.29 mm) was magnified to visualize the annotations. For (B) CellProfiler, individual colored lines denote the worm models attempting to fit the animal bodies. For (C) NemaSize, blue points denote the identified heads or tails, yellow lines denote the segmented body contours, and green lines denote the constructed skeletons. (D and E) Ivermectin dose response of the six *C. briggsae* strains processed from (D) CellProfiler and (E) NemaSize. The error bars represent one standard deviation above and below the mean.

Dose response model estimation was conducted using previously published methods [40,58]. In brief, a four-parameter log-logistic dose-response model was fitted to normalized animal lengths measured across ivermectin doses for each *C. briggsae* strain (Fig. 5D and E). For each strain-specific dose-response model, we estimated the slope (*b*) and effective concentrations producing 10%, 50%, and 90% of the maximal effect, denoted as EC_10_, EC_50_, and EC_90_, respectively.

## Results

### NemaSize achieved robust animal identification and segmentation

Before body size could be measured, NemaSize needed to identify individual animals in full-well images and segment their body masks. Errors in either step could affect downstream size measurements. Therefore, we first evaluated whether NemaSize could robustly identify and segment individual animals in the validation dataset (Table 1). We first trained YOLO26-WF to identify and segment worms in full-well images. YOLO26-WF achieved strong identification performance, with both precision and recall higher than 0.95 and an mAP50-95 of 0.859 (Table 1). However, its segmentation performance was suboptimal, with an mAP50-95 of 0.665 (Table 1 and Fig. 2B). We reasoned that the limitation likely was caused by the low number of pixels representing each individual animal at the training resolution of full-well images, which made details of body boundaries difficult to learn (see Training of YOLO26 models). The limitation of YOLO26-WF motivated us to train a second model, YOLO26-WS, using ROI images of individual worms to improve segmentation performance (see Training of YOLO26 models). YOLO26-WS achieved strong identification performance, with both precision and recall higher than 0.97 and an mAP50-95 of 0.932 (Table 1). More importantly, YOLO26-WS outperformed YOLO26-WF across all segmentation metrics, including a 0.15 increase in mAP50-95 (Table 1, Fig. 2B, and D). Together, the performance metrics supported a multiscale workflow for nematode segmentation, in which YOLO26-WF first identifies animals in full-well images, generates ROI images for individual animals, and YOLO26-WS then performs high-resolution segmentation for the animals in each ROI image (see Dual-network inference for nematode segmentation).

### NemaSize performed robust body length measurements across diverse postures

After showing that NemaSize reliably identifies and segments nematodes, we next asked how robustly NemaSize can measure animal sizes for different postures based on the segmented body masks. First, we used the validation dataset (N = 639 animals) to quantify how NemaSize measurements were consistent with the human ground-truth measurements across three merged posture classes, including straight and isolated animals, bent or curly animals, and overlapping animals (see Grouping of posture classes for data visualization and analysis; Fig. 4A). NemaSize measurements of body length showed high consistency with the human ground truth, with coefficients of determination (r²) greater than 0.98 for all three posture groups (Fig. 4A). By contrast, CellProfiler showed strong agreement with the ground truth for straight and isolated animals (r² = 0.945), but weaker agreement for bent or curly animals and overlapping animals (r² = 0.654 and 0.755, respectively) (Fig. 4A). The consistency analysis shows that both tools measured straight and isolated animals accurately, but NemaSize was in stronger agreement with human measurements for curly or overlapping animals.

Next, we asked how robustly NemaSize can measure body lengths for different postures. We therefore quantified posture-specific measurement errors for NemaSize and CellProfiler across eight posture classes using the validation dataset (N = 639 animals; see Grouping of posture classes for data visualization and analysis; Fig. 4B). NemaSize showed a mean percentage error of 3.65% across all posture classes, whereas CellProfiler showed -18.10% (Fig. 4B). The largest improvements in NemaSize were observed for curly, self-overlapping, and tiny worms, for which CellProfiler underestimated body length with median errors of -49.14%, -49.59%, and -21.26%, respectively (Fig. 4B). In comparison, NemaSize produced much smaller median errors for these three categories, including 0.97%, 1.18%, and 6.06%, respectively (Fig. 4B). The posture-specific analysis of measurement error showed that NemaSize can accurately quantify body length across diverse postures and provided more robust measurements than CellProfiler, especially for nematodes with complex postures, such as strong curvature and self overlap. The improvement reflects the importance of topology-aware skeleton extraction (Fig. 3). By classifying body topologies before skeleton extraction, NemaSize can resolve curly and self-overlapping postures according to the body geometry (Fig. 3). By contrast, methods based on model fitting, such as those used by CellProfiler [19,20], are more vulnerable to complex body geometries. The capability of NemaSize to resolve complex postures directly translates into more usable data points per imaging trial, increasing the statistical power of phenotypic screens. Even in conditions optimized for classic size measurement tools, for example straight animals, NemaSize measurements showed lower variability (SD = 3.1%) compared to CellProfiler (SD = 10.4%) (Fig. 4B). Lower measurement variability can improve the sensitivity to detect subtle differences between groups, which might be difficult to observe using classic tools or might require much larger sample sizes.

### NemaSize generate robust quantification of body width across diverse postures

In addition to body length, body width is an important morphological trait in nematodes. For example, increased body width to length ratio can distinguish Dumpy mutants from the wild type [60], whereas a decreased width to length ratio is indicative of dauer larvae [61,62]. We asked whether NemaSize could accurately measure body width across diverse nematode postures. We evaluated NemaSize width measurement error across eight posture classes using the validation dataset (N = 639 animals; see Grouping of posture classes for data visualization and analysis; Fig. S1). Because CellProfiler does not report width measurements, the analysis was performed only for NemaSize. NemaSize achieved a mean percentage error of 2.07% across all posture classes (Fig. S1), indicating that NemaSize can perform robust body width quantification across diverse postures.

### NemaSize performed body size measurements in high speed

After demonstrating that NemaSize can robustly quantify nematode body sizes, we asked how fast NemaSize could perform the analysis. We evaluated the computational speed of the NemaSize pipeline on a central processing unit (CPU) and a GPU (Table 2). On a CPU (Intel i7, 3.4 GHz), YOLO26-WF inference on a full-well image required 1.49 ± 0.69 seconds (mean ± SD), and YOLO26-WS inference on an ROI image required 0.70 ± 0.79 seconds (Table 2). A GPU (NVIDIA GeForce RTX 3090) used for processing reduced the YOLO26-WF inference to 0.08 ± 0.05 seconds and YOLO26-WS inference to 0.04 ± 0.02 seconds (Table 2). For the entire pipeline, NemaSize analyzed 1,000 animals in 14.11 minutes on the CPU and 2.23 minutes on the GPU (Table 2). To further increase computational throughput, we developed a Nextflow pipeline (https://github.com/AndersenLab/NemaSize-nf) capable of running multiple containerized instances of NemaSize in parallel on high-performance computing clusters using either CPU or GPU compute nodes. The high processing speed and capability of parallelization support the application of NemaSize in high-throughput analysis workflows.

**Table 2.**
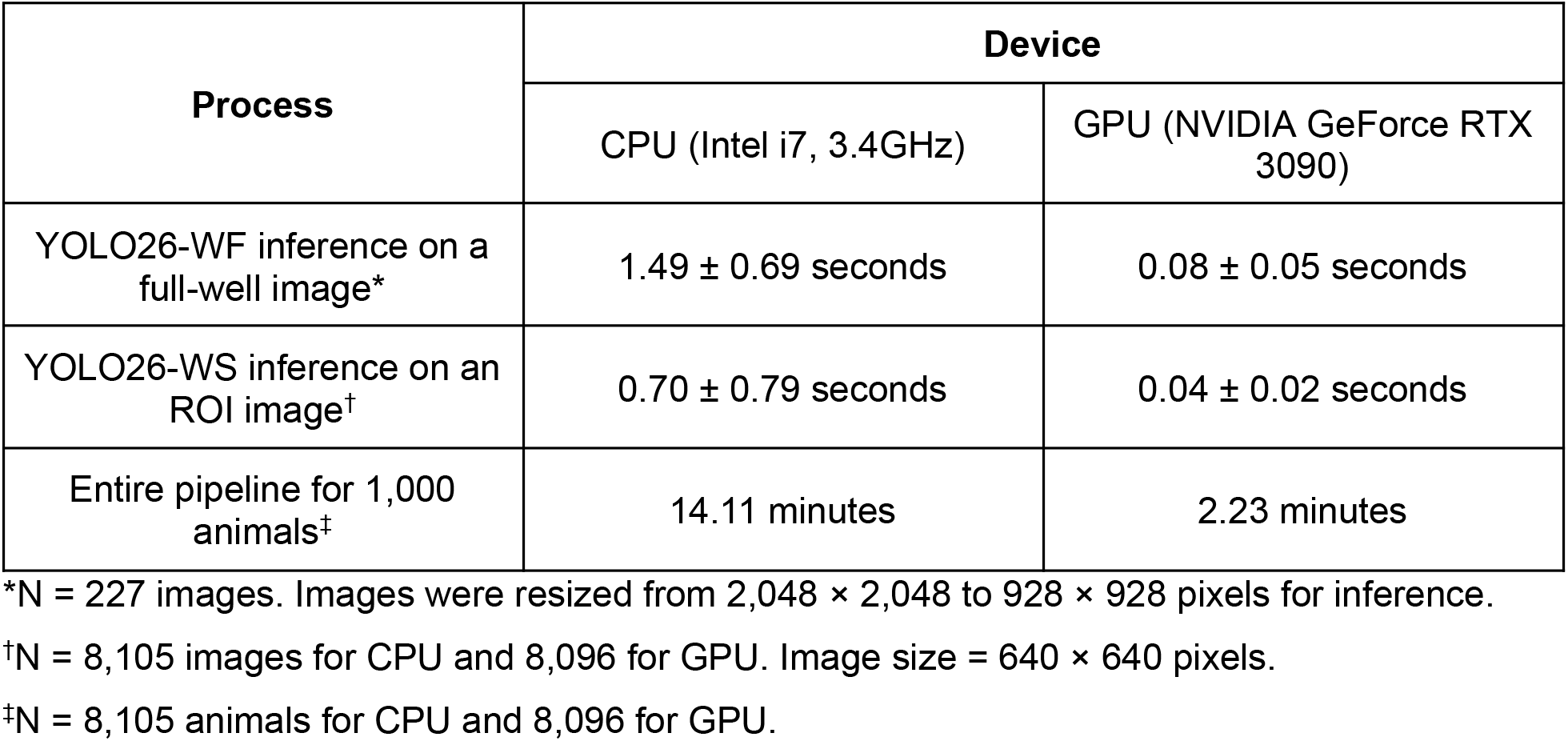
Profile of computation time for NemaSize.

### NemaSize captured accurate animal lengths and improved the interpretation of high-throughput larval development assays (HTLDA)

After demonstrating that NemaSize can accurately quantify nematode body sizes, we asked whether the improved measurement robustness could enhance the interpretation of biological data, for example larval developmental assays. We applied NemaSize to the HTLDA workflow [38–40] to quantify inhibition of *C. briggsae* larval development caused by the anthelmintic drug ivermectin (Fig. 5). We performed the analysis for six *C. briggsae* wild strains: CGC2 [63], ECA2666, ED3102, JU3237, NIC1667, and VX34 (Fig. 5A).

In zero-dose control wells (Fig. 5A-C), L1 larvae developed to the L4 stage, approximately 700 µm long [64], 48 hours after feeding (see Ivermectin dose-dependent larval development assay for *C. briggsae*). Because most animals showed curly postures (Fig. 5B), the worm models in CellProfiler could not correctly fit to the animal bodies (Fig. 5B; see Performance comparison to CellProfiler), which led to underestimation of body length across all six strains (Fig. 5A and B). In comparison, NemaSize correctly resolved the curly postures and extracted the skeleton lines (Fig. 5C), generating more accurate body length measurements (Fig. 5A). Next, we used both CellProfiler and NemaSize to quantify dose-dependent developmental inhibition in individual *C. briggsae* strains induced by ivermectin (Fig. 5D and E). We measured the normalized animal length, which represents the differences between the body length measured at a given dose and the body length measured in the zero-dose control condition. A more negative normalized animal length indicates stronger developmental inhibition (see Ivermectin dose-dependent larval development assay for *C. briggsae*). In CellProfiler measurements (Fig. 5D), all strains showed less than 100 µm reduction in body length compared to the control for doses up to 0.93 nM. The normalized animal lengths then decreased moderately as the ivermectin concentration increased to 30 nM. By contrast, NemaSize measurements better captured the dose-dependent inhibition of larval development (Fig. 5E). For all six strains, the normalized animal lengths decreased as ivermectin concentration increased. Importantly, NemaSize revealed a sharp decrease in the normalized animal length between 0.46 and 0.93 nM (Fig. 5E), which was not observed in the CellProfiler measurements (Fig. 5D). Using the dose-response curves generated by NemaSize, we obtained important estimators [40,58] to quantify the drug responses for each strain, including the slope and effective concentrations producing 10%, 50%, and 90% of the maximal effect (EC_10_, EC_50_, and EC_90_) (Table S3; see Ivermectin dose-dependent larval development assay for *C. briggsae*), which could not be accurately estimated from CellProfiler (Fig. 5D). In summary, NemaSize provided robust quantification of ivermectin-induced developmental inhibition in curly animals, a task that was difficult for CellProfiler. The more robust measurements led to more accurate estimation of drug response and improved the interpretations of HTLDA results.

## Discussion

In this study, we developed NemaSize, an AI-aided pipeline for robust body size quantification for *Caenorhabditis* nematodes. A key advantage of NemaSize is the capability to measure animals with complex postures, such as strong body curvature and self overlap, which remain challenging for other size measurement tools, including the WormToolBox of CellProfiler [19,20]. Because NemaSize can measure body sizes in animals with complex postures, it facilitates experiments that were previously difficult. For example, NemaSize could be used to quantify larval development in mutants with coiled postures [65] and to assess how different drug treatments affect development. Different from other size measurement tools that treat complex postures as technical challenges, NemaSize enables posture complexity to be analyzed as a biologically meaningful trait. For example, NemaSize could be used to study the relationship between body morphology and posture complexity, by testing whether variation in body length, width, or aspect ratio influences the probability of forming strong curvature, self overlap, or other complex postures. Because NemaSize can be adapted to high-throughput screening, it will be possible to identify mutants or wild strains with differences in body size or posture separately and also the relationship between the two traits. Although NemaSize currently processes static images, given the fast computational speed (Table 2), the pipeline could be adapted to time-lapse imaging and longitudinal tracking. For example, NemaSize could quantify the posture transitions over the course of development in individual animals, providing insights into how body morphology and postures are coordinated during development.

It is important to acknowledge the limitations of NemaSize. First, some mutually overlapping animals can produce ambiguous or incomplete body masks when the overlapping body segments are nearly collinear (Fig. S2). The problematic masks can affect downstream topological analysis and skeleton extraction. To filter unreliable skeletons, NemaSize performed a quality check during the topology classification and flagged the skeletons from unrecognized topologies. Better performance could be achieved by training the segmentation models on larger datasets containing more examples of collinear overlap. Second, NemaSize measures two-dimensional (2D) projected body size from the image rather than resolving the actual three-dimensional (3D) body structure. Although rare, the 2D measurement could be problematic when the animal forms stacked loops. Future studies could apply 3D posture estimation to further improve the accuracy of body size measurement. Third, the current YOLO26 models were trained on data collected from a single imaging platform, the ImageXpress Nano system (see Curation of annotated image dataset). Although we intentionally augmented the training images by adding variations in brightness, translation, and scaling to improve model robustness (see Training of YOLO26 models), we did not systematically test model performance on other imaging platforms. Imaging conditions with substantially different illumination, magnification, or background might require transfer learning, using the models described in this work as the initial weights. After retraining, the models could be adapted to analyze crawling animals on agar plates or other nematode species. Future collaborative data annotation and model training could further improve generalizability across imaging platforms and laboratories. Despite the limitations, NemaSize is a useful tool for robust quantification of nematode body sizes and will support broad applications in genetic and pharmacological research.

Beyond nematode body size measurement, NemaSize demonstrates a general computational strategy for analyzing elongated biological objects. Many elongated objects must be identified in a large FOV, but accurate analysis depends on boundaries or skeletons delineated at fine scales. For example, in the *C. elegans* nervous system, PVD neurons form branched dendritic arbors that extend throughout much of the body, whereas individual dendritic processes are only 35-60 nm in diameter [41–44]. Neuronal processes also form crossings and overlaps within the network [41]. Multiscale analysis could improve segmentation of fine neuronal processes, and topological classification could help identify endpoints, junctions, and loops, facilitating studies of neuronal development, regeneration, or degeneration. Similarly analysis strategies could also be applied to other elongated biological structures, such as vascular networks [45] and plant root architectures [46,47]. To support reproducible and accessible deployment of NemaSize by the community, we released the source code, pretrained YOLO26 models, and a containerized version of the pipeline (see Data availability).

## Data availability

The NemaSize source code, pretrained YOLO26 models, instructions for containerized deployment, and replication packages, including data files and analysis scripts, are available at https://github.com/AndersenLab/NemaSize. The Nextflow pipeline for parallelization is available at https://github.com/AndersenLab/NemaSize-nf.

## Acknowledgments

We would like to thank members of the Andersen laboratory for their helpful suggestions and feedback.

## Supporting information

**Figure S1.**
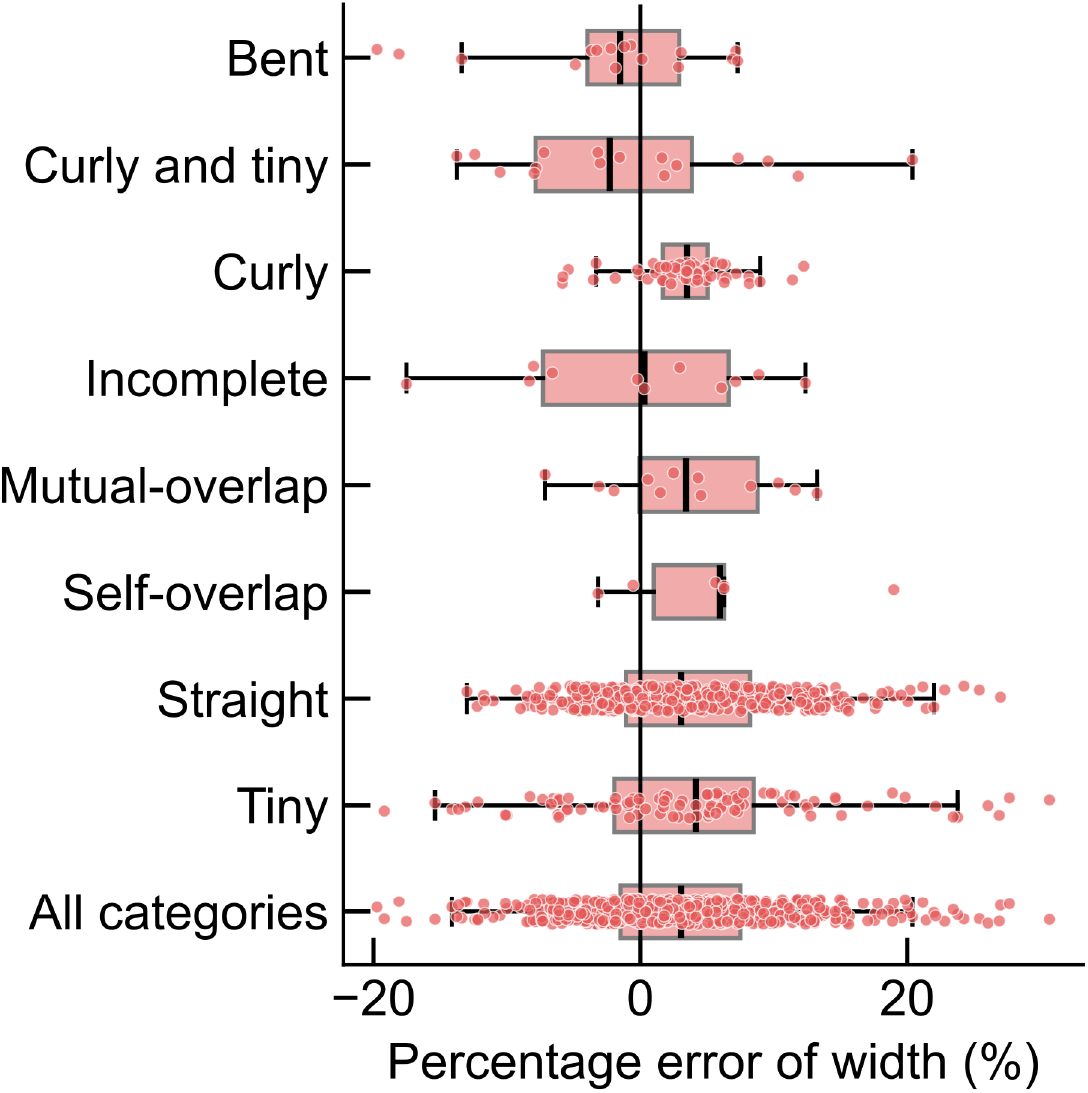
Percentage error of body width measurement for NemaSize. Each datapoint represents a single animal in individual posture classes (Tables S1 and S2). N = 16 animals for bent worms, N = 16 for curly and tiny worms, N = 60 for curly worms, N = 11 for incomplete worms, N = 12 for mutually overlapping worms, N = 6 for self-overlapping worms, N = 423 for straight worms, N = 95 for tiny worms, and total N = 639 for all categories. Data are shown as Tukey box plots with the median as a solid black vertical line, the left and right edges of the box representing the 25^th^ and 75^th^ percentiles, respectively. Whiskers extend to the most extreme data point within 1.5 times the interquartile range from the box edges.

**Figure S2.**
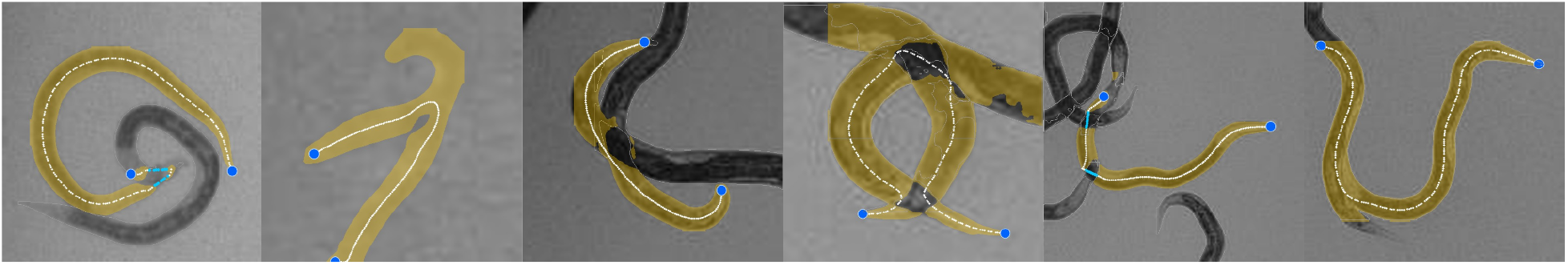
Representative failure cases for NemaSize. Each annotated ROI image depicts an example of failed skeleton extraction because of problematic mask segmentations. Yellow masks denote the segmented body masks, blue points denote the identified heads or tails, and white lines denote the constructed skeletons. From left to right, square FOV side lengths are 451 µm, 191 µm, 349 µm, 237 µm, 550 µm, and 487 µm.

**Table S1.** Definitions of posture descriptors.

| **Posture descriptor** | **Definition** |
| --- | --- |
| Bent | An animal with a single sharp bend or gradual body curvature |
| Curly | An animal with multiple sharp bends |
| Incomplete | An animal whose body is partially invisible, mainly because of occlusion by surrounding objects |
| Mutually overlapping | An animal that overlaps with at least one neighboring animal and the entire body is visible |
| Self-overlapping | An animal that overlaps with itself and the entire body is visible |
| Straight | An animal without sharp bends or noticeable body curvature |
| Tiny | An animal with a body length less than 500 $\mu\text{m}$ |

**Table S2.** Number of annotated animals for individual posture classes.

| Posture class | Count |
| --- | --- |
| Bent | 157 |
| Curly and tiny | 200 |
| Curly | 704 |
| Incomplete and curly | 42 |
| Incomplete and tiny | 70 |
| Incomplete | 296 |
| Mutually overlapping, curly, and tiny | 3 |
| Mutually overlapping and curly | 89 |
| Mutually overlapping and incomplete | 39 |
| Mutually overlapping and tiny | 25 |
| Mutually overlapping | 519 |
| Self-overlapping and tiny | 16 |
| Self-overlapping | 86 |
| Straight | 4,713 |
| Tiny and bent | 71 |
| Tiny | 930 |
| **Total** | **7,960** |

**Table S3.** Strain-specific estimators for ivermectin dose response in *C. briggsae*.

| Method | Strain | Slope* | EC<sub>10</sub> (μM)* | EC<sub>50</sub> (μM)* | EC<sub>90</sub> (μM)* |
| --- | --- | --- | --- | --- | --- |
| CellProfiler | CGC2 | 3.56e-01 ± 3.29e-02 | 1.55e-03 ± 3.80e-04 | 7.47e-01 ± 3.05e-01 | 3.60e+02 ± 3.46e+02 |
|  | ECA2666 | 4.44e-01 ± 3.65e-02 | 1.71e-03 ± 3.57e-04 | 2.40e-01 ± 6.15e-02 | 3.37e+01 ± 2.18e+01 |
|  | ED3102 | 2.97e-01 ± 2.89e-02 | 1.19e-03 ± 3.38e-04 | 1.92e+00 ± 1.03e+00 | 3.09e+03 ± 3.81e+03 |
|  | JU3237 | 3.93e-01 ± 3.57e-02 | 1.80e-03 ± 4.08e-04 | 4.84e-01 ± 1.72e-01 | 1.30e+02 ± 1.11e+02 |
|  | NIC1667 | 3.37e-01 ± 3.36e-02 | 1.83e-03 ± 4.70e-04 | 1.25e+00 ± 6.10e-01 | 8.52e+02 ± 9.57e+02 |
|  | VX34 | 3.72e-01 ± 3.31e-02 | 1.27e-03 ± 3.09e-04 | 4.66e-01 ± 1.68e-01 | 1.70e+02 ± 1.48e+02 |
| NemaSize | CGC2 | 4.55e-01 ± 2.15e-02 | 4.24e-05 ± 9.78e-06 | 5.31e-03 ± 4.85e-04 | 6.64e-01 ± 1.73e-01 |
|  | ECA2666 | 3.90e-01 ± 1.98e-02 | 2.55e-05 ± 7.00e-06 | 7.12e-03 ± 7.65e-04 | 1.99e+00 ± 6.64e-01 |
|  | ED3102 | 4.56e-01 ± 2.13e-02 | 4.39e-05 ± 1.02e-05 | 5.44e-03 ± 4.91e-04 | 6.75e-01 ± 1.72e-01 |
|  | JU3237 | 4.36e-01 ± 2.13e-02 | 3.87e-05 ± 9.43e-06 | 5.96e-03 ± 5.80e-04 | 9.18e-01 ± 2.60e-01 |
|  | NIC1667 | 4.08e-01 ± 1.99e-02 | 2.92e-05 ± 7.46e-06 | 6.37e-03 ± 6.58e-04 | 1.39e+00 ± 4.27e-01 |
|  | VX34 | 5.38e-01 ± 2.42e-02 | 7.27e-05 ± 1.45e-05 | 4.31e-03 ± 3.41e-04 | 2.56e-01 ± 5.14e-02 |

\*Values are reported as point estimate ± standard error (SE).

